# Nanoscale distribution of nuclear sites analyzed by superresolution STED-ICCS

**DOI:** 10.1101/753228

**Authors:** M. Oneto, L. Scipioni, M.J. Sarmento, I. Cainero, S. Pelicci, L. Furia, P.G. Pelicci, G.I. Dellino, P. Bianchini, M. Faretta, E. Gratton, A. Diaspro, L. Lanzanò

## Abstract

Deciphering the spatiotemporal coordination between nuclear functions is important to understand its role in the maintenance of human genome. In this context, superresolution microscopy has gained considerable interest as it can be used to probe the spatial organization of functional sites in intact single cell nuclei in the 20-250 nm range. Among the methods that quantify colocalization from multicolor images, image cross-correlation spectroscopy (ICCS) offers several advantages, namely it does not require a pre-segmentation of the image into objects and can be used to detect dynamic interactions. However, the combination of ICCS with super-resolution microscopy has not been explored yet.

Here we combine dual color stimulated emission depletion (STED) nanoscopy with ICCS (STED-ICCS) to quantify the nanoscale distribution of functional nuclear sites. We show that STED-ICCS provides not only a value of colocalized fraction but also the characteristic distances associated to correlated nuclear sites. As a validation, we quantify the nanoscale spatial distribution of three different pairs of functional nuclear sites in MCF10A cells. As expected, transcription foci and a transcriptionally repressive histone marker (H3K9me3) are not correlated. Conversely, nascent DNA replication foci and the Proliferating cell nuclear antigen (PCNA) protein have a high level of proximity and are correlated at a nanometer distance which is close to the limit of our experimental approach. Finally, transcription foci are found at a distance of 130 nm from replication foci, indicating a spatial segregation at the nanoscale. Overall, our data demonstrate that STED-ICCS can be a powerful tool for the analysis of nanoscale distribution of functional sites in the nucleus.

**Statement of significance:** Several methods are available to quantify the proximity of two labeled molecules from dual color images. Among them, image cross-correlation spectroscopy (ICCS) is attractive as it does not require a pre-segmentation of the image into objects and can be used to detect dynamic interactions. Here, we combine for the first time ICCS with superresolution stimulated emission depletion (STED) microscopy (STED-ICCS) to quantify the spatial distribution of functional sites in the nucleus. Our results show that STED-ICCS, in addition to quantifying the colocalized fraction, detects characteristic nanometer distances associated to correlated nuclear sites. This work shows that STED-ICCS can be a powerful tool to quantify the nanoscale distribution of functional sites in the nucleus.

## Introdution

Healthy genome regulation and maintenance rely on the proper spatiotemporal coordination between several nuclear functions. Alterations in chromatin organization are often linked to human diseases (1). An example is represented by DNA transcription and replication, two fundamental genomic processes that can potentially compete for the same DNA template. Replication and transcription both occur within the highly packed chromatin environment and must be tightly coordinated in time and space to avoid interference and generation of DNA damage (2). Alterations in the coordination of replication and transcription represent a major source of genomic instability, a hallmark of cancer (3). Therefore, deciphering the coordination between nuclear functions at a high spatial and temporal resolution is important to understand its role in health and disease. Genome-scale sequencing methods provide high-resolution maps of spatiotemporal regulation of genomic processes (4, 5). However, they do not provide any information on spatial localization and lack single cell resolution. In this context, optical imaging methods are emerging as complementary tools to investigate genome organization and structure in intact single cell nuclei (6).

The simplest approach to study spatial organization of two labeled molecules consists in the analysis of dual color images for the presence of colocalized signal, i.e. signal that overlaps within the two detection channels. Colocalization is a measurement of the co-distribution of two probes at a spatial scale defined by the resolution of the optical microscope. In conventional optical microscopy, this spatial scale is limited by diffraction to about 250 nm. For this reason, alternative strategies have been developed to investigate cellular processes at the nanoscale. For instance, a popular method to probe molecular distances in the nanometer range is Forster Resonance Energy Transfer (FRET) (7–9) but, unfortunately, FRET is not sensitive when distances are larger than about 10 nm. Similarly, *in situ* Proximity Ligation Assay (PLA) is a powerful method to visualize proximity of two labeled species but its sensitivity does not exceed 40 nm (10, 11). This scenario changed dramatically with the introduction of superresolution microscopy (also called nanoscopy), namely the ensemble of microscopy techniques providing optical resolution below the diffraction limit (12). For instance, in single molecule localization microscopy (SMLM), the fluorophores are sequentially switched on and off and localized with high precision on each frame, resulting in images with a resolution down to ~20 nm (13–15). In stimulated emission depletion (STED) microscopy the fluorophores are selectively switched off at the periphery of the diffraction-limited detection spot by a second, doughnut-shaped laser beam, producing an immediate improvement of spatial resolution down to ~50 nm (16). Thus, superresolution can be used to probe spatial organization in the 20-250 nm range, which corresponds to the range of higher order organization of chromatin in the nucleus. This nanoscale spatial organization can be studied in a ‘static’ way, from images of fixed cells or in a ‘dynamic’ way, from acquisition of superresolution data in live cells.

Methods that provide a quantitative measure of colocalization from static multicolor superresolution images can be divided in two major groups: object-based methods, which perform a segmentation of the image into objects prior to analyzing their relative spatial distributions, and pixel-based methods, which extract correlation coefficients from pixel intensities (17, 18). In general, object-based analysis can be performed on any type of superresolution image as long as the target objects are well resolved and identified. In particular, object-based methods are often the methods of choice in SMLM, where the acquired data are already segmented into a list of x y coordinates of individual molecules (19–23). In principle, object-based analysis provides a full description of the spatial distribution of the two labeled species: in fact, knowledge of the objects coordinates allows (i) mapping the locations of the specimen with a higher level of proximity and (ii) performing a statistical analysis of the relative distance between the particles. On the other hand, a great advantage of pixel-based methods is that they do not require a pre-segmentation of the images into objects but rely on the calculation of coefficients from the pixel intensity values (24, 25). Pixel-based methods are routinely applied to quantify spatial distribution in multicolor superresolution images (26–28).

A method to study interactions in a ‘dynamic’ way is Fluorescence Cross-Correlation Spectroscopy (FCCS), the dual channel version of Fluorescence Correlation Spectroscopy (FCS) (29). In FCCS, the fraction of interacting particles is extracted from the analysis of temporal intensity fluctuations originating from changes in fluorophore concentration within a small observation volume, typically defined by the focal spot of a confocal microscope (30, 31). Both FCS and FCCS have been applied to study the mobility and interactions of molecules in the nucleus (32–34). Interestingly, the very same formalism of FCS and FCCS can be applied to the analysis of the spatial intensity fluctuations found in images. Image Correlation Spectroscopy (ICS) and Image Cross-Correlation Spectroscopy (ICCS) are the spatial variants of FCS and FCCS, respectively (35). Notably, ICCS can be used as a pixel-based method to analyze the spatial distribution in static images as well (36). Thus, ICCS appears as an extremely versatile method that can offer several advantages compared to other analysis approaches, namely it does not require a pre-segmentation of the image into objects and can be used to detect dynamic interactions (37–39). However, despite this potential, the combination of ICCS with superresolution microscopy has not been fully explored yet.

Here, we show that the combination of dual color STED nanoscopy with ICCS (STED-ICCS) can be used to quantify the relative nanoscale spatial distribution of distinct nuclear foci. In particular, we show that STED-ICCS can provide, to some extent, some of the attractive features of an object-based analysis but without requiring a pre-segmentation of the super-resolved image. In fact, we are able to (i) map the locations within nuclei with a higher level of proximity and (ii) determine if the particles are correlated at a certain nanoscale distance. First, the analysis is tested on simulated images of nuclear foci at variable density and compared with an object-based analysis performed on the same simulated datasets. Then, the STED-ICCS analysis is tested on dual color STED images of model samples based on DNA origami bearing green and red fluorophores located at a characteristic distance of 20 nm or 100 nm. These data demonstrate that STED-ICCS can provide nanoscale information on the distance between correlated particles. Finally, to validate the STED-ICCS method on a biological sample, we quantify the relative nanoscale spatial distribution of three different pairs of nuclear sites in MCF10A cells. As expected, transcription foci and a transcriptionally repressive histone marker (H3K9me3) show the minimum level of colocalization and random relative distance distribution. Conversely, nascent DNA replication foci and the Proliferating cell nuclear antigen (PCNA) protein have a high level of proximity and are correlated at a nanometer distance which is close to the limit of our experimental approach. Notably, nascent DNA replication foci and transcription foci are found to be partially correlated but at a distance of ~100 nm, indicating that the two functional sites are spatially segregated at the nanoscale. Overall, these data demonstrate that STED-ICCS can be a powerful tool for the analysis of relative nanoscale spatial distribution of functional sites in the nucleus.

## Results

### Simulations: STED-ICCS is a robust method to characterize relative spatial distribution at high densities of foci

To compare STED-ICCS with object-based analysis, we simulated dual color images of nuclear foci at variable densities (Fig.1). In each channel, the nuclear foci were simulated as *N* point-like particles distributed in a 16 µm wide circular region (40) and convoluted with a Gaussian Point Spread Function (PSF) with a Full Width Half Maximum (FWHM) of 3 pixels, corresponding to 120 nm. We simulated distributions of foci randomly distributed in each channel (uncorrelated), with a fraction *f*_coloc_=0.25 of foci colocalizing in the two channels (colocalized) and with a fraction *f*_coloc_=0.25 of foci colocalizing only in a specific sub-portion of the image (colocalized in a zone). The total number of foci in each channel was varied from *N*=100 to *N*=10000. These simulations are intended to test the analysis workflow and not as a model of the more complex distributions and patterns that can be found in real samples.

**Fig.1.**
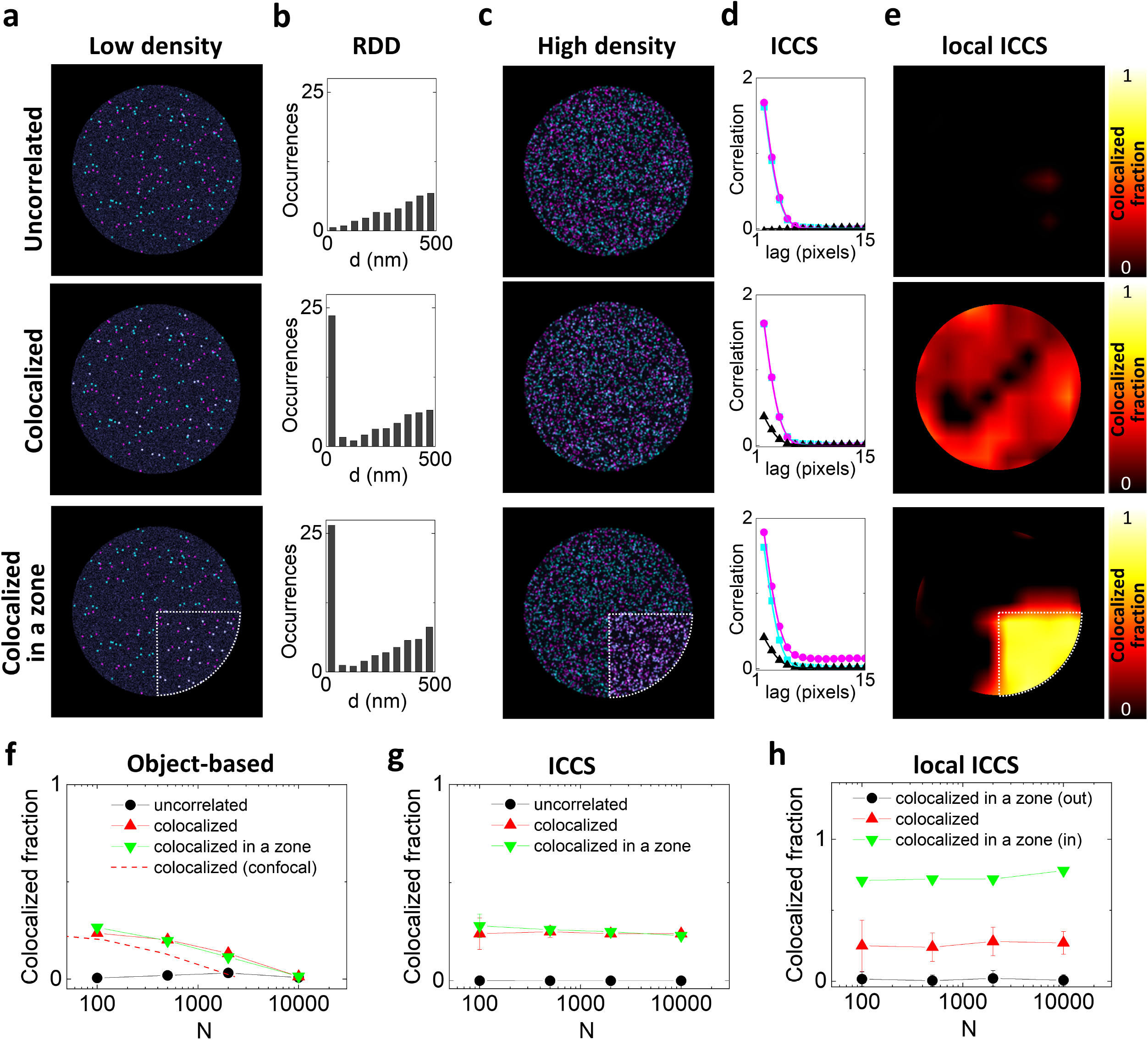
Comparison between STED-ICCS and object-based analysis on simulated data. (a) Simulated dual color images of nuclear foci at low density (total number of foci, *N*=100), assuming a random distribution of foci in each channel (uncorrelated), a 25% of foci colocalization in the two channels (colocalized) and a 25% of foci colocalizing only in a specific sub-portion of the image (colocalized in a zone), indicated by the dashed contour. The data were simulated with a PSF size of FWHM=120 nm. (b) Relative distance distributions (RDD) obtained by object-based analysis of the simulated images. (c) Simulated dual color images of nuclear foci at high density (total number of foci, *N*=2000). (d) Spatial correlation functions of the images shown in (c). Shown are the cross-correlation function (black triangles) and the single channel auto-correlation functions (magenta dots and cyan squares) along with the corresponding fits (solid lines). (e) Local ICCS maps of the images shown in (c). The colormap represents the value of parameter *f*_ICCS_ calculated on a moving subregion of 69×69 pixels. (f) Colocalized fraction extracted from object-based analysis of the simulated data as a function of the number of foci *N*. The dashed red line shows the trend for the colocalized sample when the PSF size is FWHM=250nm. (g) Colocalized fraction extracted by ICCS analysis of the simulated data as a function of the number of foci. (h) The value of *f*_ICCS_ extracted from the local ICCS map inside (in) and outside (out) the colocalization zone for the ‘colocalized in a zone’ sample, compared with the value of *f*_ICCS_ in the ‘colocalized’ sample.

At low density, the foci are easily localized in the XY plane and the relative distance between every center in one channel and every center in the other channel, can be calculated from their center positions (Fig. 1a) according to the localization-based approach. The resulting relative distance distribution (RDD) has a linearly growing trend for the uncorrelated sample, as expected from purely geometrical considerations (Fig.1b). For the other two samples in Fig.1a (colocalized and colocalized in a zone), the RDD shows an additional peak at a distance zero, corresponding to the fraction of colocalized foci, superimposed with the linear trend due to the random distribution (Fig.1b). At higher densities of foci (Fig.1c), the colocalized fraction retrieved with the object-based analysis (Fig.1f, triangles) deviates from the simulated value (*f*_coloc_=0.25). In particular, a deviation of 20% from the expected value is obtained when the number of foci is *N*=500 (Fig.1f). For comparison, considering a FWHM value typical of diffraction-limited microscopy, FWHM_conf_=250 nm, a deviation of 20% is obtained already when the number of foci is *N*=115. This is due to the lower accuracy in determining the center positions of the foci during the segmentation process.

On the other hand, ICCS is able to determine the colocalized fraction independently of the density of foci (Fig.1g). In ICCS, the colocalized fraction *f*_ICCS_ is extracted from the fitting of the image auto- and cross-correlation functions (41, 42). The amplitude of the cross-correlation function is zero for the uncorrelated sample (Fig.1d, top), whilst is positive for both colocalized samples (Fig.1d, mid and bottom). In the range of foci densities explored here, the value of colocalized fraction retrieved by ICCS is consistent with the simulated value (Fig.1g), in keeping with previous studies (36). Thus, STED-ICCS can be used to determine the fraction of cross-correlated particles also when the density of foci is too high for non-super-resolved methods or, equivalently, when the improvement of spatial resolution is not sufficient to resolve the foci.

An apparent disadvantage of ICCS, in comparison with object-based approaches, is that it provides only an average description of the properties of the sample in the region analyzed. In fact, the two correlated samples shown in Fig.1c (colocalized and colocalized in a zone) produce almost identical correlation functions (Fig.1d) despite a very different colocalization pattern. Thus, ICCS can be used to extract a value of the colocalized fraction at high density of foci but does not specify where the foci are colocalized. To partially overcome this limitation, we performed local ICCS analysis, i.e. ICCS performed on small sub-regions of the image (Fig.1e, h). It has been previously shown that, if the spatial correlation function is calculated locally, one can get maps of physical parameters such as protein velocity (43), particle size (44) or diffusion coefficient of a probe (45, 46). Local ICCS can be used to generate a map of the value of the colocalized fraction extracted by fitting the local auto- and cross-correlation functions (Fig.1e). The local ICCS maps show that the two correlated samples (colocalized and colocalized in a zone) have a very different pattern, despite containing the same total fraction of colocalized foci. The spatial resolution of the local ICCS map depends on the size of the sub-image employed for the local analysis. For instance, in the example of Fig.1e, the size of the sub-image was set to 69 pixels.

### Measurements on model systems: STED-ICCS detects particles correlated at a distance

Another limitation of STED-ICCS, when compared to object-based analysis, is that it cannot be used to perform a complete statistical analysis of the relative distance between the particles. This feature of object-based analysis is particularly useful to detect characteristic nanometer distances associated to inter-molecular complexes (20). Here we aim to show that, even if it is not able to provide a full statistical analysis, STED-ICCS can provide information on the average distance between correlated particles.

In STED-ICCS, the shape of the cross-correlation function is dependent upon the distance *d* between the two probes and the effective spatial resolution of the STED microscope in the two channels. For two single spots located at distance *d* along a given direction, if the single channel 2D auto-correlation functions have width *w*_11_ and *w*_22_, the corresponding 2D cross-correlation function has a width equal to *w*_cc_=((*w*_11_^2^+*w*_22_^2^)/2)^1/2^ and is shifted from the origin of an amount *d* along the same direction (47). In most real cases, however, the particles are found at all possible orientations with respect to each other. This has two effects on the cross-correlation function, namely a reduction of its amplitude and an increase of the value of its width compared to the value *w*_cc_, expected for perfectly coaligned spots. The broadening of the cross-correlation function can be evaluated as Δ*w*=*w*_12_−*w*_cc_. Importantly, Δ*w* is a parameter sensitive only to the average distance between correlated particles, whilst the amplitude-related parameter *f_ICCS_* is sensitive also to the relative amount of correlated particles. Thus, the Δ*w* can be used to estimate the distance associated to the correlated particles. To actually convert values of Δ*w* into values of distance, we performed simulations of particles correlated at variable distances and measured the broadening of the cross-correlation function (Supplementary Fig. S1).

To demonstrate this effect, we performed STED-ICCS on optical nanorulers, i.e. DNA origami structures designed to contain the same two fluorophores used in our experiments (Chromeo-488 and Atto-532) at a well-defined distance of either *d*=20nm or *d*=100 (Fig.2a, b). First, we characterized the samples by object-based analysis. The spatial resolution achieved in these STED images was ~95 nm in the green channel, and ~130 nm in the red channel, as determined from the FWHM of line profiles (Supplementary Fig. S2). Considering an average number of photons detected per spot per channel *N*~30 and a 10% background noise level, this translates to an estimated localization precision σ_G_~13 nm and σ_R_~20 nm in the green and red channels, respectively (48). This propagates to an expected uncertainty (σ_G_^2^+σ_R_^2^)^1/2^~24 nm in the estimation of the relative distance *d*. The distributions of distances obtained from the object-based analysis of STED images of optical nanorulers are reported in (Fig.2c, d). They contain a peak superimposed to a linear growing component. The distance peak value found in the relative distance distribution (RDD) of the 100-nm nanorulers was d_100nm_=100±25 nm (mean±s.d.), in keeping with the expected value. These data indicate that, at the imaging conditions of this sample, we can measure distances with an accuracy σ_d_~25 nm. This sets a lower limit to the distances that can be detected with our experimental setup. In line with this, the distance peak value found in the RDD of the 20-nm nanorulers was d_20nm_=40±30 nm.

**Fig.2.**
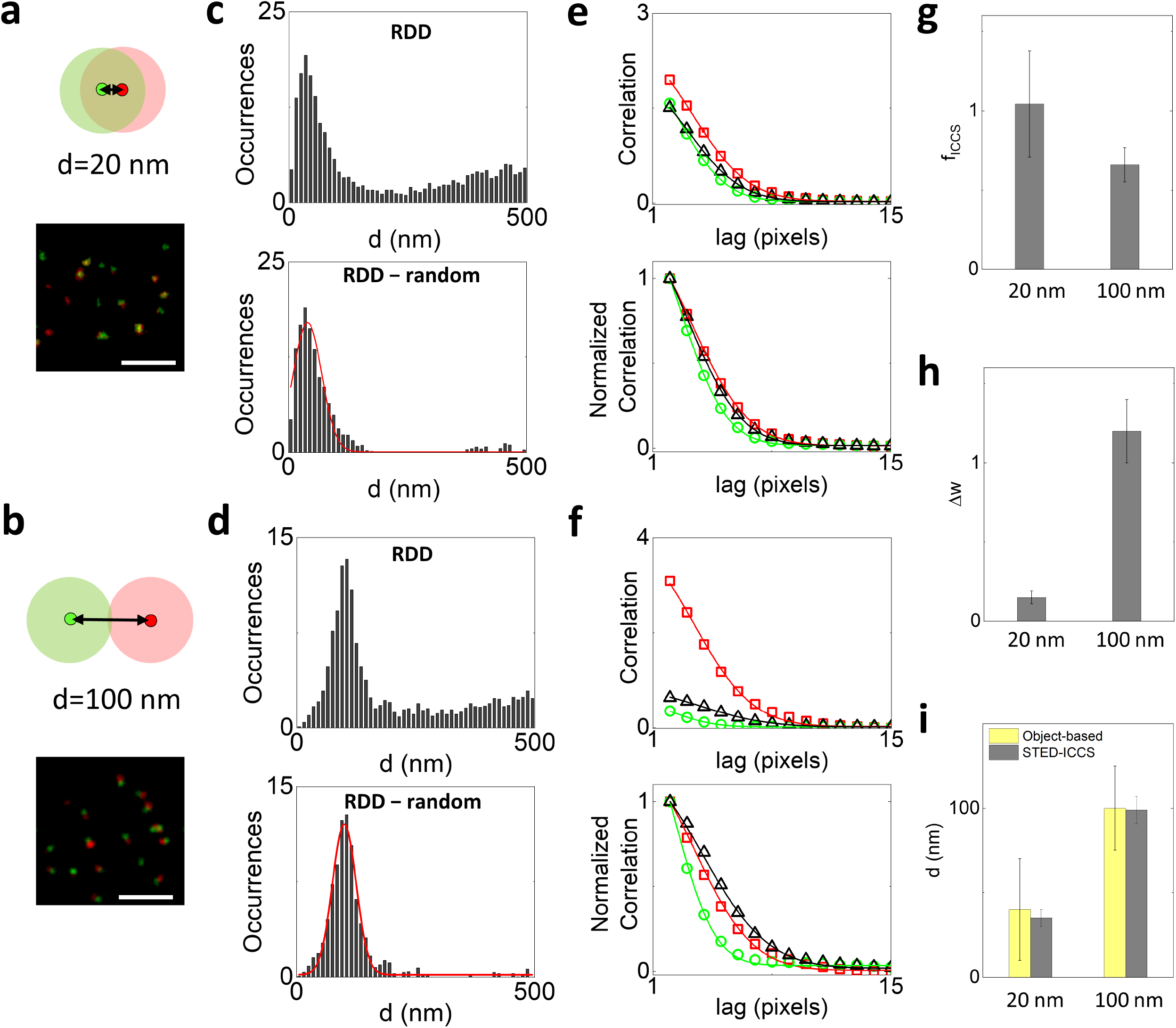
STED-ICCS of model samples. (a,b) Schematic drawing and representative dual color STED images of the optical nanorulers. These model systems consist of Chromeo-488 and Atto-532 fluorophores located at a fixed distance of 20 nm (a) and 100 nm (b). The shaded circles schematically represent the PSF. Scale bar 1 µm. (c,d) Object-based analysis for the 20-nm (c) and 100-nm (d) nanorulers. Shown are the relative distance distribution histograms (RDD) before (top) and after (bottom) subtraction of the uncorrelated random component. Solid red lines are Gaussian fits of the data (d_20nm_=40±30 nm, d_100nm_=100±25 nm, mean±s.d.). (e,f) Raw (top) and normalized (bottom) correlation functions of the representative images shown in (a,b). Shown are the cross-correlation function (black triangles) and the red (red square) and green (green circles) channel auto-correlation functions along with the corresponding fits (solid lines). (g) Colocalized fraction extracted from STED-ICCS analysis. (h) Cross-correlation function broadening obtained from STED-ICCS. (i) Values of distances determined by object-based analysis and STED-ICCS.

Then we performed STED-ICCS on the same samples. As expected, STED-ICCS detected a positive cross-correlation for both samples (Fig.2e, f), with the colocalized fraction being higher in the 20-nm nanorulers (*f*_ICCS_=1.04±0.3, mean±s.d.) than in the 100-nm nanorulers (*f*_ICCS_=0.66±0.11, mean±s.d.) (Fig.2g). More interestingly, the parameter Δ*w*, related to the broadening of the cross-correlation function (Fig.2h), was higher in the 100-nm nanorulers (Δ*w*=1.2±0.2 pixels, mean±s.d.) than in the 20-nm ones (Δ*w*=0.15±0.04 pixels, mean±s.d.). According to the simulations, the measured values of Δ*w* correspond to the distance values *d*_ICCS_=99±8 nm, for the 100-nm nanorulers, and *d*_ICCS_=35±5 nm, for the 20-nm nanorulers, which are in agreement with the peak values found with object-based analysis (Fig.2i). Note that the smaller uncertainty in the value of *d*_ICCS_ is due to the fact that each STED-ICCS measurement is performed on an image containing several tens of nanorulers whilst, in the object-based analysis, the distance is measured on each identified pair of objects.

Thus, the relative broadening of the STED-ICCS cross-correlation function can provide nanoscale information on the distance between particles that are not distributed randomly.

### Measurements in cell nuclei: STED−ICCS quantifies the relative nanoscale distribution of functional nuclear sites

As validation of our STED-ICCS method, we characterized the nanoscale spatial distribution of three different pairs of functional nuclear sites in the diploid mammary epithelium cells MCF10A. We compared the nanoscale spatial distribution of i) transcription foci (labeled through incorporation of the nucleotide analogue BrU) *versus* a transcriptionally repressive histone marker (H3K9me3) (Fig.3a), ii) nascent DNA replication foci (labeled through incorporation of the nucleotide analogue EdU) *versus* the replication machinery protein proliferating cell nuclear antigen (PCNA), during early S phase (Fig.3b), iii) transcription foci (BrU) *versus* nascent DNA replication foci (EdU), during early S phase (Fig.3c). The STED-ICCS analysis was compared with an object-based analysis performed on the same dataset.

**Fig.3.**
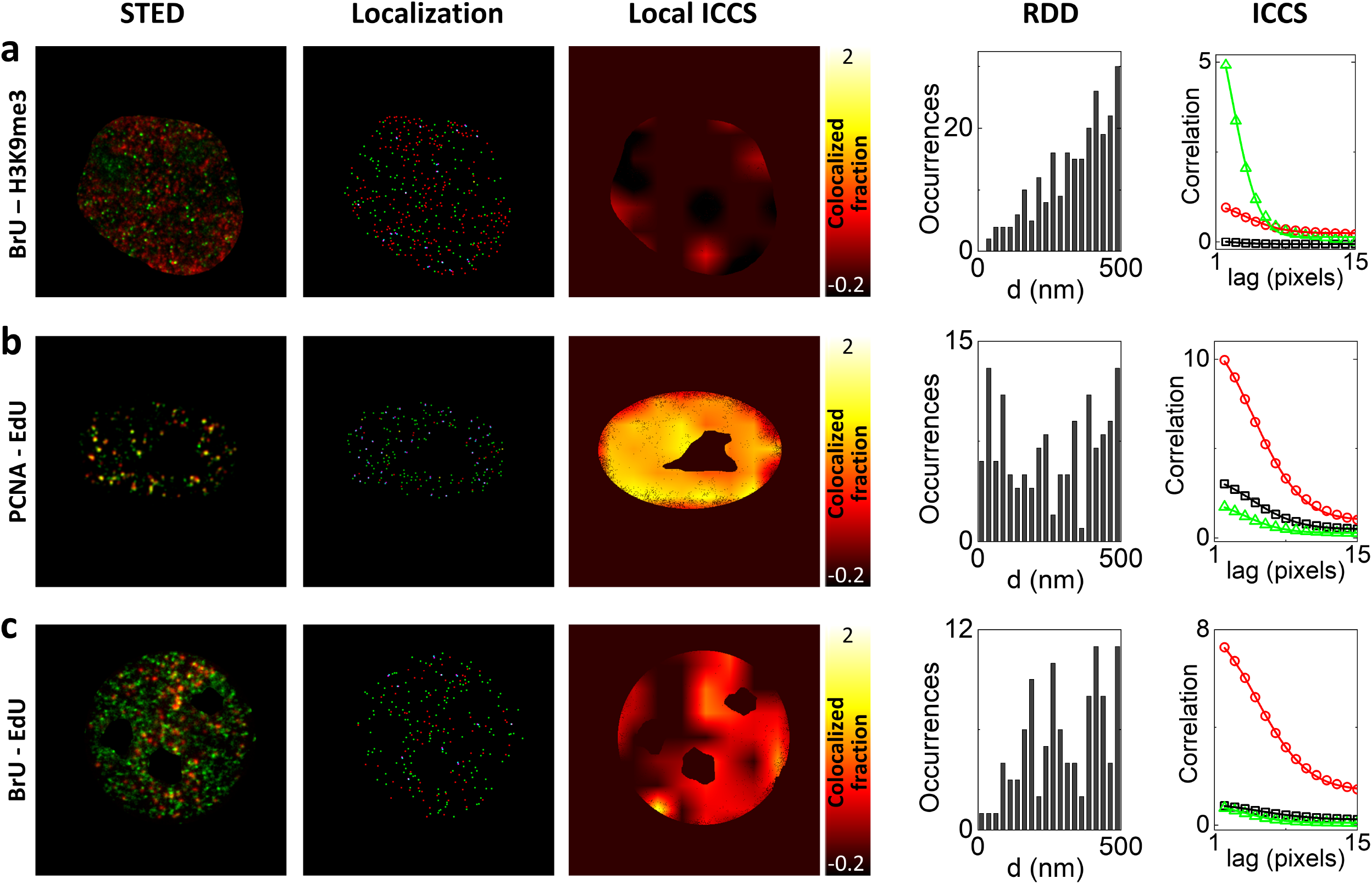
Analysis of nuclear foci by STED-ICCS and object-based localization. Analysis of representative STED images of MCF10A cells acquired upon labelling of (a) BrU (green) and H3K9me3 (red), (b) PCNA (green) and EdU (red), (c) BrU (green) and EdU (red). Shown are (from left to right) the dual color STED image, the positions of the colocalized (cyan) and non-colocalized (red and green) foci recovered by particle localization, the map of the colocalized fraction recovered by local ICCS, the RDD histogram and the spatial correlation functions recovered by ICCS. The ICCS plot shows the cross-correlation function (black triangles) and the red (red squares) and green (green circles) channel auto-correlation functions along with the corresponding fits (solid lines).

Representative dual color STED images of the three samples are reported in Fig.3. For each image are shown the map of localized spots along with the calculated RDD, the image auto- and cross-correlation functions and the map of the colocalized fraction obtained by local ICCS (Fig.3). Overall, there was an agreement between the colocalized fraction extracted by the object-based analysis, *f_obj_*, and that extracted by STED-ICCS, *f_ICCS_*, for the different samples (Supplementary Fig. S3). Differences in the absolute values (*f_ICCS_* was systematically higher than *f_obj_*) are probably related to the specific settings of each analysis. For instance, in the object-based analysis, an intensity threshold value was set manually for each image whilst, in the ICCS analysis, we did not perform any background subtraction. Notably, the local ICCS maps can be used to visualize, in STED-ICCS, the regions of the nuclei with a higher level of proximity, similarly to what can be done with object-based analysis (Fig.3). The average co-localization fractions retrieved from the STED-ICCS analysis are reported in Fig.4c. Transcription foci and the transcriptionally repressive histone marker H3K9me3 show the minimum level of colocalization (*f_ICCS_*=0.02±0.05, mean±s.d., n=17 cells), in keeping with the evidence that H3K9me3 is an epigenetic marker of repressive heterochromatin, replicating during late-S phase and not associated with active transcription (49). This result is confirmed by object-based analysis, where transcription and H3K9me3 show a random-like RDD (Fig.4a, b). Conversely, DNA replication foci and the PCNA protein have the maximum level of colocalization (*f_ICCS_*=1.01±0.16, mean±s.d., n=19 cells), in agreement with the role of the PCNA protein in orchestrating the replication process (50). Transcription foci and DNA replication foci in early-S phase cells exhibit an intermediate level of colocalization (*f_ICCS_*=0.36±0.09, mean±s.d., n=22 cells). However, this analysis does not show if this difference in the level of colocalization is due to a different distribution at the nanoscale.

**Fig.4.**
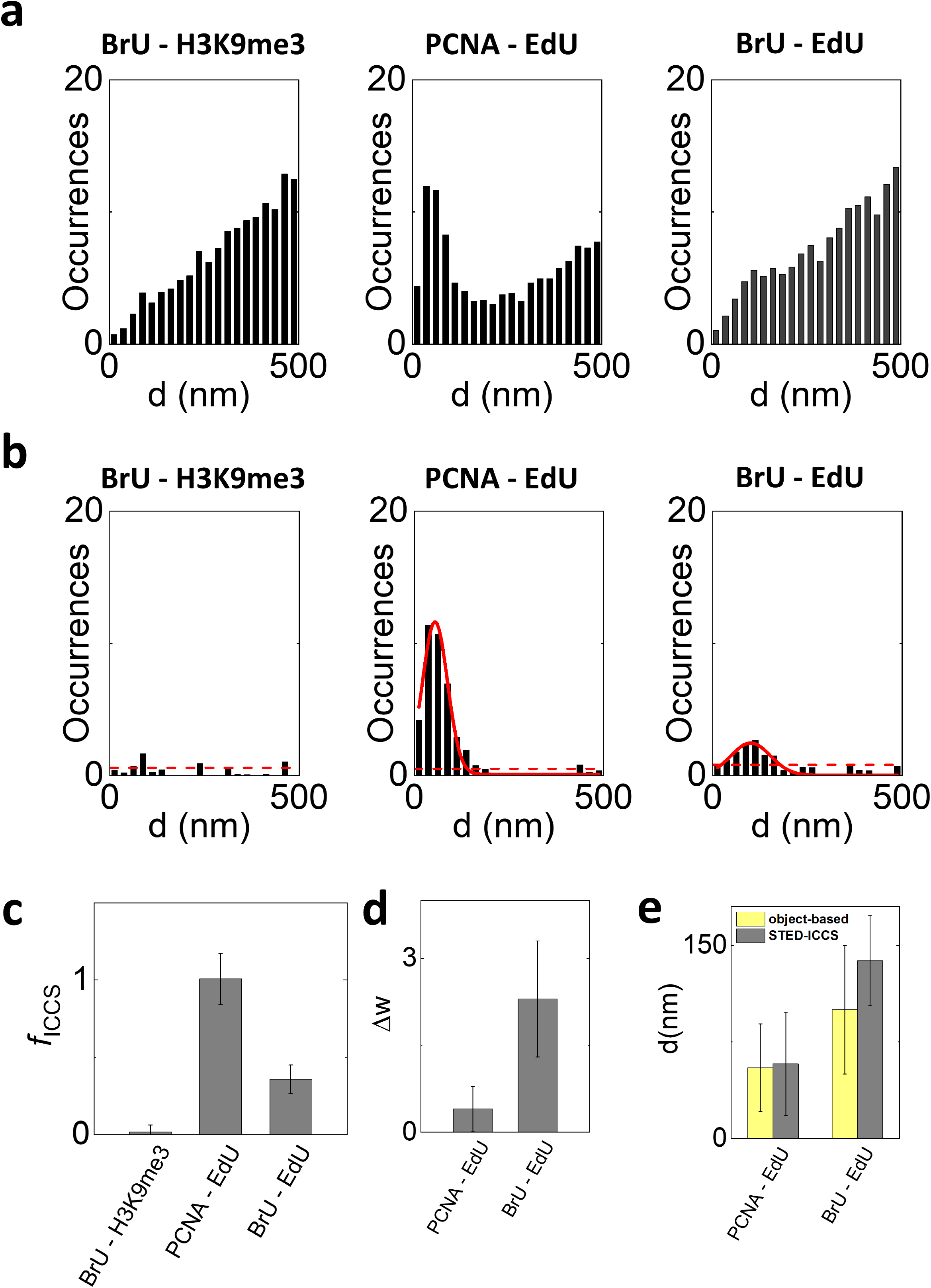
Relative nanoscale spatial distribution of the investigated nuclear sites in MCF10A cells. (a,b) Cumulative results of the object-based analysis. Shown are the relative distance distribution histograms (RDD) before (a) and after (b) subtraction of the uncorrelated random component. Solid red lines are Gaussian fits of the data (PCNA-EdU: d=55±34 nm, mean±s.d., n=19 cells; BrU-EdU: d=100±50 nm, mean±s.d., n=22 cells). The dashed red lines indicate the standard deviation of the data in the range 250-500 nm after subtraction of a linear fit of the uncorrelated component. (c,d) Cumulative results of the STED-ICCS analysis. (c) Colocalized fraction extracted from STED-ICCS analysis. (d) Cross-correlation function broadening extracted from STED-ICCS. (e) Values of distances extracted by object-based analysis and STED-ICCS for the correlated samples.

To get insight on the relative nanoscale distribution of the correlated functional sites, we analyzed the broadening of the cross-correlation function, Δ*w* (Fig.4d), and compared it with the relative distance distribution (RDD) derived from the object-based analysis (Fig.4a, b). The RDD of DNA replication foci and the PCNA protein revealed a peak distance of 55 nm (d_PCNA-EdU_=55±34 nm, mean±s.d., n=19 cells). STED-ICCS detected a broadening Δ*w*_PCNA-EdU_=0.4±0.4 pixels (mean±s.d., n=19 cells) corresponding to a distance value *d*_PCNA-EdU_=58±40 nm. The values of distance detected by the two methods are very close to the limit of our experimental approach, as determined with the 20-nm optical nanorulers (*d_obj_*=40 nm and *d*_ICCS_=35nm). A higher level of inaccuracy might also come from the use of primary and secondary antibodies. Moreover, we analyzed only single optical sections that represent only a projection of the distribution of nuclear foci in three dimensions. This could result in underestimation of the recovered distance values. Besides these technical considerations, we should also take into account that the EdU signal labels the DNA sequences that were replicating during the previous 20 minutes and that PCNA, not only tethers polymerases to DNA for during replication, but also participates in non-replicative DNA synthesis events, such as those occurring during DNA repair, and other cell functions that extend well beyond DNA synthesis.

The RDD of transcription and DNA replication foci in early-S phase cells exhibited a quasi-random relative distance distribution (Fig.4b). After subtraction of the random component, the RDD showed a small peak indicating a positive correlation at a distance of d_BrU-EdU_=100±50 nm (mean±s.d., n=22 cells) (Fig.4b). Interestingly, STED-ICCS detected a broadening Δ*w*_BrU-EdU_=2.3±1.0 pixels (mean±s.d., n=22 cells) corresponding to a distance value *d*_BrU-EdU_=138±35 nm (Fig.4b, d, e). Thus, STED-ICCS was able to reveal a correlation between these nuclear sites in the ~100-nm range, similarly to the object-based analysis. The non-random distribution in the BrU-EdU sample is consistent with the cell population which was analyzed; during early-S phase, replicating DNA is mainly euchromatic and gene-rich. Most of the genes contained within these regions are generally expressed during early S-phase, and importantly, although their transcription is time-controlled in order to avoid transcription-replication clashes, it is still very close to ongoing replication, not only in space, but also in time, both prior and after DNA synthesis. Thus, in early-S phase, transcription and replication operate on the same portion of the genome, thus explaining proximity of the corresponding BrU and EdU signals (51). In summary, STED-ICCS detected a small but positive correlation between replication and transcription (quantified by the parameter *f_ICCS_*) but indicated spatial segregation at the nanoscale (quantified by the parameter *d_ICCS_*), in keeping with a long-standing model of genome organization suggesting spatial segregation between replication and transcription (2, 51).

## Discussion

In this work, we have explored the use of super-resolution STED imaging, combined with image cross-correlation spectroscopy (ICCS), to investigate the relative nanoscale spatial distribution of nuclear foci. There are three main technical points that characterize STED-ICCS. The first important aspect is that, being based on the calculation and fit of the image spatial correlation functions, STED-ICCS does not require a pre-segmentation of the images into objects. To illustrate the consequences of this point, we have performed ICCS and object-based analysis on simulated data of point-like particles at different density/spatial resolution. These simulations indicated that ICCS could be applied even at high densities of foci, whilst the object-based analysis was affected by a decreasing accuracy in the pre-segmentation process when the spatial resolution was not sufficient to resolve the foci. A second technical aspect is that in STED-ICCS the colocalized fraction is estimated from the amplitude of the image cross- and auto-correlation functions, calculated over the image. Compared to object-based analysis, this calculation does not specify where the foci are colocalized. In this respect, we have shown that a local STED-ICCS analysis can be used to map the value of the colocalization coefficient across the sample and partially compensate for this limitation. The third important aspect is related to the detection of characteristic correlation distances. In the object-based approaches, it is possible to perform a complete statistical analysis of the relative distance between the particles and detect characteristic nanometer distances associated to inter-molecular complexes (20). In this respect, we have shown that the broadening of cross-correlation function in STED-ICCS can also provide quantitative information on the nanoscale distance between correlated particles.

To validate our approach, we performed STED-ICCS on dual color STED images of model and biological samples and compared the results with an object-based analysis performed on the same datasets. In particular, the object-based quantification was presented in terms of a relative distance distribution (RDD), representing the histogram the distance values between particles. However, we expect that any other type of quantitative analysis applied to the list of coordinates of the positions of the foci, such as radial distribution functions or Ripley’s functions (20, 21), would give similar results. We quantified by STED-ICCS the relative nanoscale distribution of three pairs of functional nuclear sites. Notably, STED-ICCS was able to detect not only a value of colocalized fraction but also the characteristic correlation distance associated to correlated nuclear sites. In particular, PCNA was found in close association with EdU-labeled replication foci, with a detected distance of ~50 nm, very close to the experimental limit of the analysis. On the other hand, transcription foci were found at a distance of 130 nm from EdU-labeled replication foci, indicating a spatial segregation at the nanoscale despite both transcription and replication taking place on the same portion on the genome during the early S phase of the cell cycle. Overall, there was a good agreement between STED-ICCS and the object-based analysis, demonstrating that, even without requiring a pre-segmentation, STED-ICCS can provide some of the most attractive features of an object-based analysis.

An important point to discuss is how general is the applicability of our method. We have shown experimental data obtained on model and fixed biological samples by dual color STED imaging. In particular, in the conditions of our experiments, we achieved a lateral resolution in the order of ~100 nm. The same type of nanoscale ICCS analysis could be applied to other types of ‘static’ superresolution images, including images obtained with SMLM. In the case of SMLM, a segmentation of the data is already available making object-based co-localization analysis the method of choice. Nevertheless, we believe that the ICCS analysis could be useful as an independent cross-validation of the results obtained with object-based approaches. Even more interesting appears the application of our method to the analysis of ‘dynamic’ superresolution images, for which an object-based analysis is less straightforward. In our data, a spatial resolution of the order of ~100 nm was sufficient to characterize the relative nanoscale spatial distribution of the three pairs of functional nuclear sites investigated in our samples. The same spatial resolution could be achieved, in principle, with many live-cell superresolution imaging approaches. For instance, Structured illumination Microscopy (SIM), and its point-scanning equivalent Image Scanning Microscopy (ISM), are superresolution techniques compatible with live cell imaging, even if their resolution improvement is limited to a factor of ~2 (12). The same STED nanoscopy includes several variants developed to reduce the STED beam intensity and its potentially photo-damaging effects (40, 52, 53). Thus, we expect that our ICCS formalism could be applied to live cell super-resolved images obtained, for instance, by dual color STED- or SIM-based setups. In this perspective, we believe that our work could be useful to establish, in the near future, a new type of dynamic analyses that cannot be obtained in other static superresolution techniques.

## Materials and Methods

### Cell culture

Human mammary epithelial cells MCF10A were grown in DMEM (Merck KGaA, Darmstadt, Germany):Ham’s F12K (Thermofisher Scientific, Waltham, MA, USA) medium (1:1) containing 5% horse serum, 1% penicillin/streptomycin, 2 mM L-glutamine, 10 μg/mL insulin, 0.5 μg/mL hydrocortisone, 50 ng/mL cholera toxin (all from Merck KGaA) and 20 ng/mL EGF (PeproTech, Rocky Hill, NJ, USA), at 37 °C in 5% CO_2_. For fluorescence microscopy measurements, cells were seeded on glass coverslips coated with 0.5% (w/v) gelatin (Sigma Chemical Co.) and cultured for 18h in growth medium. Cells were incubated for 20 min with the synthetic nucleotides 5-ethynyl-2’- deoxyuridine 10 μM, EdU (Thermofisher Scientific) and 5-Bromouridine 10 mM, BrU (Sigma Chemical Co.) to allow labelling of replication and transcription, respectively. Upon nucleotide incorporation, cells were washed with PBS and fixed with 4% paraformaldehyde (w/v) for 10 min at room temperature.

### Sample preparation

For immunostaining, MCF10A cells were permeabilized with 0.1% (v/v) Triton X-100 in blocking bugger (BB), composed of 5% w/v bovine serum albumin (BSA) in PBS, for 1h at room temperature. To recognize nascent DNA (transcription) and RNA (replication) filaments, incorporated BrU and EdU were labeled, respectively. For BrU detection, cells were incubated overnight at 4 °C with a primary rabbit antibody anti-BrdU (Rockland Immunochemicals Inc., Limerick, PA, USA) diluted at 1:1000 in BB, followed by three 15 min rinses in BB. EdU incorporation was detected using the Click-iT EdU imaging kit (Thermo Fisher Scientific) according to the manufacturer instructions but replacing the kit’s azide-Alexa488 with Azide-PEG3-biotin-conjugated (Merck) at a 1:500 dilution to allow the subsequent immunostaining. After the click reaction, cells were washed with PBS and incubated with the primary mouse antibody anti-biotin (Merck) diluted at 1:1000 in BB, for 1h at room temperature. To detect PCNA and EdU, incorporated EDU nucleotides were conjugated with biotin-azide using the Click-iT EdU imaging kit as previously described. Samples were then incubated with primary antibodies anti-biotin (1:1000 dilution in BB) and anti-PCNA in rabbit IgG (Santa Cruz Biotechnology Inc. Dallas, Texas, USA) (1:500 dilution in BB) at the same time for 1h at room temperature. To detect transcriptionally active and repressed chromatin, we labeled BrU and Histone 3 lysine 9 tri-methylated (H3K9me3), respectively. The cells were simultaneously incubated with primary antibody anti-BrdU as previously described and anti-H3K9me3 in mouse IgG (Abcam, Cambridge, UK) at a 1:800 dilution, overnight at 4 °C in BB.

After three 15 min rinses in BB, cells were incubated with secondary antibodies (1:200 dilution) for 1h at room temperature in BB, upon which cells were washed with BB three times for 15 min. The secondary antibodies were Atto532 and Chromeo488 conjugated with goat anti-mouse IgG (Rockland Immunochemicals Inc.) and anti-rabbit IgG (Abcam, Cambridge, UK), respectively. All samples were washed with PBS and water before mounting with Mowiol (Sigma).

DNA origami structures containing the two fluorophores Chromeo488 and Atto532 at a well-defined distance of either d=20 nm or d=100 nm were purchased from GATTAquant (custom nanorulers, GATTAquant, Germany).

### Experiments

All imaging experiments were performed on a Leica TCS SP5 gated-STED microscope, using an HCX PL APO 100x 100/1.40/0.70 Oil immersion objective lens (Leica Microsystems, Mannheim, Germany). Emission depletion was accomplished with a 592 nm STED laser. Excitation was provided by a white laser at the desired wavelength for each sample. For imaging of MCF10A nuclei, Chromeo488 was excited at 470 nm and its fluorescence emission detected at 480-530 nm, with 1-10 ns time gating using a Hybrid detector (Leica Microsystem). Atto532 excitation was performed at 532 nm and the emission collected between 545-580 nm by a Hybrid detector, with time gating of 2.5-6 ns. The two channels were acquired in line-sequential mode and the excitation and depletion power were adjusted separately for each channel. 512×512 pixel images were acquired with a pixel size of 40 nm. Similar settings were used for imaging the 100 nm and 20 nm nanorulers.

### Simulations

Dual color images of nuclear foci were simulated using MATLAB. For each channel, the object consisted in a variable number *N* of point-like emitters distributed randomly inside a circular area with a diameter of 16 µm (40). The images were made by 512×512 pixels with a pixel size of 40 nm. The maximum total number of photons detected from a single pixel position from a single particle was set to S=40. For each channel, the object was convolved with a gaussian Point Spread Function (PSF) with Full Width Half Maximum (FWHM) of 120 nm and a uniform background level B=3 was added within the circular region. Finally, the resulting images were corrupted by Poisson noise. To simulate the uncorrelated sample, the foci were distributed randomly in both channels. To simulate a partially colocalized sample, a number *N*/4 of foci in the second channel was set to have the same coordinates of *N*/4 foci in the first channel. To simulate the sample colocalized in a zone, the colocalized foci were forced to be in a specific quarter of the circular area. The total number of foci in each channel was varied from *N*=100 to *N*=10000.

### Image cross-correlation spectroscopy (ICCS) and local ICCS

The ICCS analysis was performed in MATLAB using a custom code. The 2D image correlation functions were calculated as:

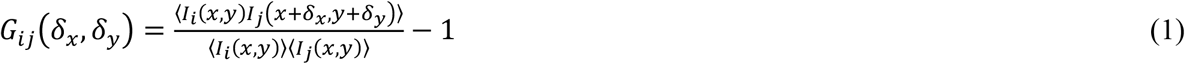

where *I*_1_(*x*,*y*) and *I*_2_(*x*,*y*) are the images in the first and the second channel, respectively, and the angle brackets indicate averaging over all the selected pixels of the image. The two autocorrelation functions were obtained by setting i=j=1 and i=j=2, respectively, whereas the cross-correlation function was obtained by setting i=1 and j=2. The numerator in Eq.(1) was calculated by a 2D-FFT (Fast Fourier Transform) algorithm. Before calculation, a Region of Interest (ROI) corresponding to the nucleus was defined and all the pixels outside this ROI were assigned an intensity value equal to the average value inside ROI, as reported previously (41). This step is useful to minimize the effects of nuclear borders on the correlation functions.

The 2D correlation functions were then converted into radial 1D correlation functions *G_ij_*(*δ_r_*) by performing an angular mean, as described previously (44). The resulting radial correlation functions were then fitted to a gaussian model:

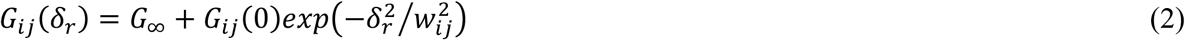

in order to extract the amplitude parameters *G_ij_*(0) and the width parameters *w_ij_*.

The amplitude parameters were used to calculate the coefficients of colocalizations M_1_ and M_2_ (36):

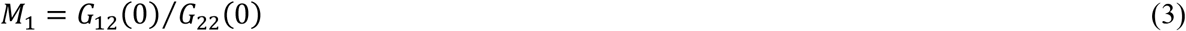

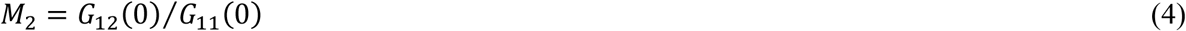

The colocalized fraction *f*_ICCS_ was then calculated as the arithmetic mean of the two coefficients *M*_1_ and *M*_2_. The broadening of the cross-correlation function with respect to the corresponding auto-correlation function has been evaluated with the parameter Δ*w*=*w*_12_−*w*_cc_, where *w*_cc_=((*w*_11_^2^+*w* ^2^)/2)^1/2^.

To discard bad cross-correlation function fits, we excluded i) fits with a chi-square value 50 times larger than the chi-square value of the auto-correlation function fit; ii) fits with a width *w*_12_ too different compared to the average autocorrelation function widths (*r_w_*<0.5 or *r_w_*>2, where *r_w_*=*w*_12_/(*w*_11_*w*_22_)^1/2^); iii) fits with a negative offset G_∞_ (G_∞_<−0.2 *G*_12_(0)). In all these cases, we set *f*_ICCS_=0.

The local ICCS analysis consisted in performing ICCS iteratively on small, 69×69 pixels wide, sub-regions of the full-size image. For each sub-region, the local auto- and cross-correlation function were calculated and the parameters *G_ij_*(0), *w_ij_* and *f*_ICCS_ were extracted as described for ICCS analysis. In addition to the conditions described above, we excluded local auto-correlation fits with an amplitude 10-times larger than the amplitude of the global autocorrelation function and local cross-correlation fits with an amplitude 2-times larger than the amplitude of the local autocorrelation function. The calculation was limited to *N*_samp_=100 sampling sub-regions centered on *N*_samp_ pixels uniformly distributed over the image. Then the resulting values were interpolated to produce a map of the same size of the full-size image.

### Object-based analysis and relative distance distribution (RDD)

The central coordinates of the foci in each channel were obtained using the JaCoP plugin (54) in ImageJ (55). For each channel, an intensity threshold was set manually. The minimum size of the particles was set to 2 pixels. The algorithm provided the coordinates of all the localized particles in each channel and the values of distance from each particle in the first channel from all the particles in the second channel. In the simulations, particles were considered colocalized if their distance was lower than 50 nm. In the biological samples, particles were considered colocalized if their distance was lower than 140 nm. All the values of distance were used to build the relative distance distribution (RDD) histogram. To estimate the random component in the cumulative RDD histogram, a linear fit of the data through the origin was performed in the range 250- 500 nm. This linear component was then subtracted from the data to obtain a RDD without the random component.

## Author Contributions

LL, AD, EG, MF, GID, PGP designed research; MO, LS, MJS, LF, PB performed research; LL and LS wrote software; LL, MO, LS, MJS, IC, SP analyzed data; LL wrote the paper with input from all the authors.

## Acknowledgements

LL, MO and MJS were supported by Fondazione Cariplo and Associazione Italiana per la Ricerca sul Cancro (AIRC) through Trideo (Transforming Ideas in Oncological Research) Grant number 17215. The research leading to these results has also received funding from AIRC under MFAG 2018 - ID. 21931 project – P.I. Lanzanò Luca.

